# Beyond Neural Noise: Critical Dynamics Predict Slower Reaction Times in Adults With and Without ADHD

**DOI:** 10.64898/2026.03.13.711705

**Authors:** Alessandra DallaVecchia, Nicolas Zink, Samantha Reina O’Connell, Samantha S. Betts, Sean Noah, Austin Hillberg, Mercedes T. Oliva, James T. McCracken, Rory C. Reid, Gregory V. Simpson, Sarah L. Karalunas, Vince D. Calhoun, Agatha Lenartowicz

## Abstract

Neural variability observed during task has frequently been interpreted as “noise,” assumed to obscure meaningful signal and thus something to be minimized both analytically by researchers and functionally by the brain. Changes to the signal-to-noise ratio have been proposed as a possible neural mechanism behind the increased reaction-time variability (RTV) in attention deficit hyperactivity disorder (ADHD). However, not all variability is the same in some cases, variability can have some underlying “statistical structure” that can be beneficial to information processing. The challenge lies in distinguishing meaningful variability from random noise. The edge-of-synchrony critical point, which describes a system poised between synchronous and asynchronous regimes, could be a useful theoretical framework to study the association between neural and behavioral variability in ADHD, as it describes how variability changes across different regimes. In this study, we investigate whether changes in criticality and oscillatory dynamics preceded slower behavioral responses during a bimodal continuous performance task in ADHD. In adults with and without ADHD, neural dynamics shifted towards criticality, and thus greater statistical structure, prior to slower responses. Adults with ADHD did exhibit increased variability across both task and passive viewing conditions, but this increased variability was associated with greater proximity to criticality compared to controls. Thus, our findings suggest that increased RTV in ADHD and during attention lapses is related to structured variability (with greater proximity to criticality) and not necessarily random “noise.” Notably, this effect was not evident at the between-subject level, underscoring the limitations of relying on between-subject correlations to infer neural mechanisms. These findings also highlight the value of capturing multiple features to distinguish between neural models.

**Impact Statement:** Reaction time variability (RTV) and increased instances of slow response times in attention deficit hyperactivity disorder (ADHD) have been hypothesized to reflect increased neural “noise” or variability. However, our findings show that this increased variability is associated with criticality, a dynamical regime linked to improved computation. Furthermore, the core difference between ADHD and controls is not overall variability, but rather that ADHD neural dynamics operate closer to this critical regime during both task and rest.

## Introduction

Behavioral variability is an increasingly important indicator of neural variability, especially in neuropsychiatric disorders. In attention-deficit hyperactivity disorder (ADHD), for example, increased behavioral variability is consistently found during sustained attention tasks as increased reaction time variability (RTV) which has been linked to a greater frequency of longer, slower reaction times (RTs) [1, 2]. Mechanistically, it has been hypothesized that increased RTV could be due to increased neural “noise” in ADHD [3, 4, 5]. Evidence of this noise arises from the presence of flatter slopes in the power spectrum of EEG data between children with and without ADHD [6, 7]. Why would flattening of the slope be indicative of noise? The power spectrum in EEG reflects the activity of oscillators in the brain, usually from 0.1 - 50 Hz and its slope represents the interactions between these oscillators. Changes in this slope can give insights into the dynamics of the system. In the waking brain, the spectral slope approximates a 1/f frequency distribution which is associated with long range temporal correlations (LRTCs). LRTCs refer to a phenomenon where a transient event does not just happen and vanish, but the effects of the event ripple across time in the system. In other words, the system has memory of prior events as their effects propagate across time. When the system has too many transient events and is unable to propagate them – losing its memory – then the spectral distribution becomes flatter, eventually resulting in a flat line or “white noise” [8]. A flattening of the slope is thus related to an increase in random fluctuations with little to no propagation, effectively like adding random “noise” in the system [9]. Possible neural mechanisms of this flattening and its connection to “noise” arise from simulation work that shows that flattening of the slope can arise with higher background rates of de-coupled firing (or asynchronous spiking) and more neural variability [10, 9]. This negative association with flattening is then further substantiated by evidence that individuals with flatter slopes both at rest and within-task show worse behavioral performance [11, 12, 6].

However, the term “noise” has a negative connotation which makes it seem like flattening is always bad. When a distribution has too steep of a slope, approaching “brown noise,” the system becomes overly correlated in time and functionally rigid [8], suggesting that sometimes flattening may be desirable. Indeed, the spectral slope is flatter in task than at rest [13, 14, 15](cf. [16]), which would mean that flattening of the slope, and thus a shift towards neural variability, is *beneficial* for cognitive processing. This contradicts the association of flatter slopes with noisier brains as predicted by the above-mentioned “noise” hypothesis. This contradiction also emphasizes how *inter-*individual (between-subject) differences in RTV and spectral slope do not always translate to *intra*-individual (within-subject) associations between RTV and neural signals like spectral slope. A possible explanation for such opposing effects is that both the individual slope at rest (between-subject differences) and the *degree*of flattening of the slope during task (a within-subject modulation) contribute to behavioral performance, suggestive of possibly distinct drivers or sources. Thus, it is essential to better understand the mechanisms that drive these changes in system dynamics and neural variability in order to distinguish between variability that is detrimental from that which is functionally meaningful.

A possible mechanism through which the brain can modulate neural variability and stability is provided by a dynamic systems framework known as the Critical Brain Hypothesis [8, 17]. This framework may help explain the associations between behavioral performance and within-versus between-subject changes by linking the spectral slope to underlying network states. The hypothesis posits that the brain operates near a “critical point” or a transition point between two dynamical regimes. While various models of criticality are theoretically plausible, one particularly relevant type is “edge-of-synchrony.” Here, the critical point is the exact boundary between an asynchronous irregular regime (where activity is highly variable and disorganized, analogous to a flat, white noise distribution) and a synchronous regime (characterized by rigid, highly reliable, but inflexible dynamics, analogous to a steep, brown noise distribution) (Fig. 1). A system can be tuned to either phase, but it is precisely at the transition between these two extremes that the balanced 1/f spectrum emerges, alongside several beneficial computational properties such as maximal dynamic range, information communication, complexity, and LRTCs [17, 18, 8, 19, 20]. Theoretically, a system poised at criticality would have access to both a high degree of variability and synchronous activity allowing for optimal information communication and sensitivity to incoming stimuli. It is similar in concept to the Yerkes-Dodson law that describes the relationship between arousal and performance as an “inverted-U,” such that there is an “optimal” level of arousal and too little or too much are associated with decrements in performance [21].

**Figure 1:**
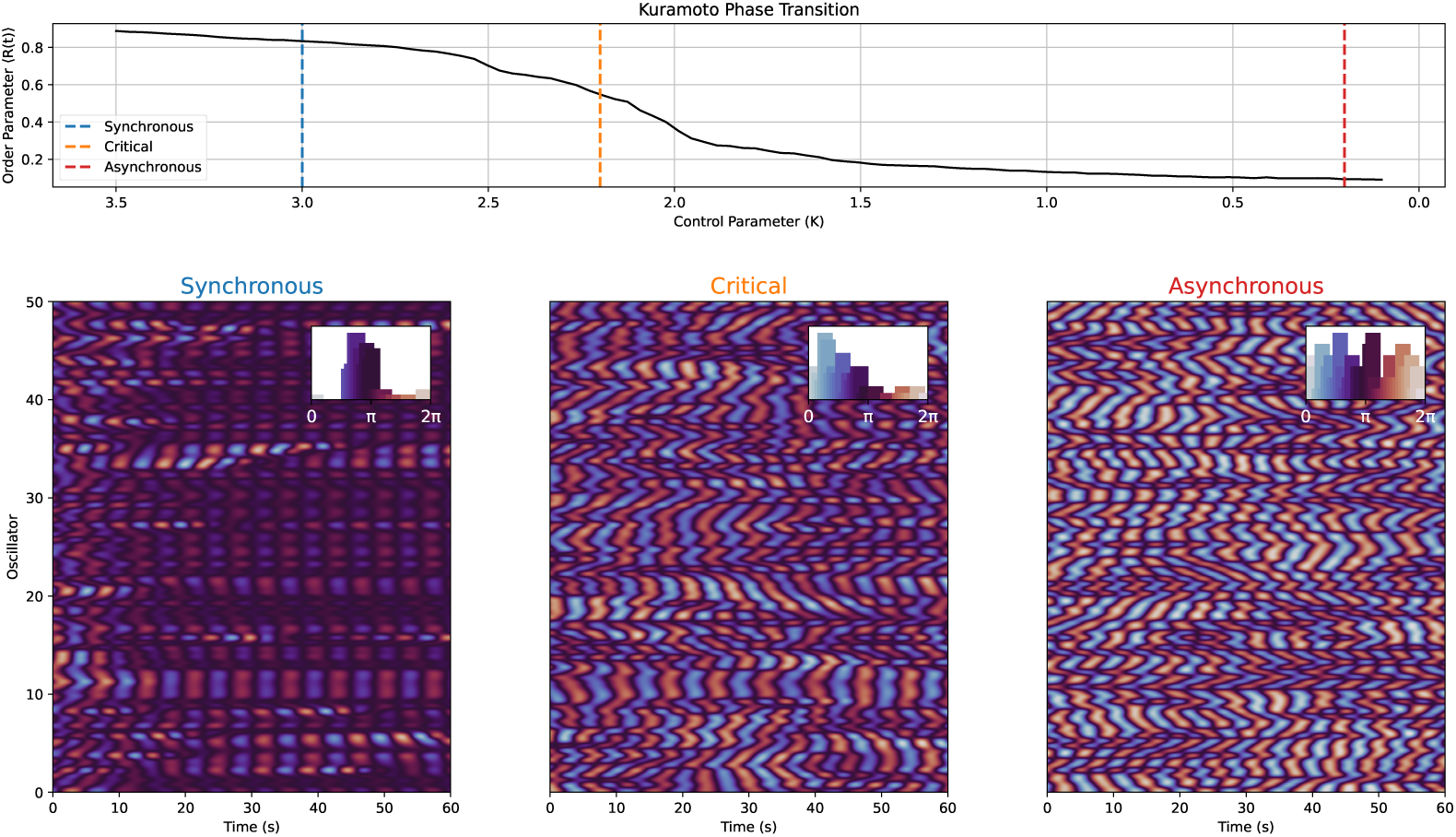
Kuramoto Model of Edge of Synchrony Criticality. This figure shows an example of how activity changes in the different regimes around the critical point called the edge-of-synchrony using a Kuramoto-Sakaguchi model [22] with an alpha (8 - 12 Hz) natural frequency and lag (*α*) of 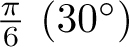 to induce a coupling delay between oscillators. In the top row, the dynamics of the model (R(t) or the order parameter) are described at different values of “K” (called the control parameter). Between 2 and 2.5 K, the dynamics start to change dramatically, transitioning from a regime of low synchrony (asynchronous) to one of high synchrony. In systems capable of more complex dynamics, like the Kuramoto-Sakaguchi and the brain, criticality or critical phenomena can emerge at a range of values and not a singular point. On the bottom row, the dynamics at three K values corresponding to points in the synchronous, critical, and asynchronous regimes are represented by raster plots with mean-centered oscillator phase activity over a period of 60 seconds. The inset axis show the phase distributions for all three regimes. In the synchronous regime, there are “islands” of oscillators that are synchronized and the phase distribution is highly concentrated around *π*. The oscillator phases in the asynchronous regime, in contrast, are varied over time and source (oscillator) as can also be seen by the distribution of phases in bar-plot on the left of the raster. In the middle, the critical regime has both characteristics from the other two regimes: there are some “islands” of synchrony, but also places and times where the oscillators are out of synchrony. The phase distribution during the critical regime has some concentration of phases (near 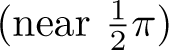 like the synchronous regime, but is not so constrained.

Differences in within-versus between-subject effects may be explained by the system shifting away and towards criticality in order to exploit characteristics of the different regimes. For example, the system may perform better at a certain task when it is in a more stable phase (for our specified phase transition, it would be the synchronous phase) as information can be transferred with more certainty. The opposite is also true – a task may require higher sensitivity if it requires more task switching so a shift towards criticality is more beneficial. Thus, the proximity to criticality may be dynamically tuned to balance between higher variability (e.g., sensitivity) and stability of responses, as a function of task demands [23, 24, 25, 26]. Viewed through a criticality lens, the flattening of the spectral slope of neural activity during task might reflect this deviation from criticality [27, 28, 29] in the direction of more excitability (towards the asynchronous phase). In fact, there is supporting evidence that during attention-focused tasks, the brain deviates further from criticality [30, 31, 32], but the direction of this deviation is unclear. Both the results from spectral slope and LRTCs would appear to suggest that task states are tuned slightly towards the asynchronous regime. However, the magnitude of this shift appears important as *within-task* worse behavioral performance has been associated with flatter slopes [6]. Therefore, although accumulating evidence suggests a link between criticality and behavioral performance, it remains uncertain how strongly deviations from criticality predict performance, and in which direction these deviations typically occur (within-subject effects). Furthermore, given the distance from criticality is theoretically a parameter that can be dynamically tuned, it is equally important to know how this modulation interacts with its “starting” parameters (between-subject effects). In other words, does the brain modulate the transition between task and rest the same way if,at rest, it starts closer or further from criticality (between-subject effects)? Resolving these questions would provide critical insight into the neural dynamics that shape both between-subject differences (e.g., does increased noise explain the differences in slope changes between individuals with ADHD vs. controls) and within-subject changes in neural dynamics during task and rest.

To address this unknown, in the current study we looked at several overlapping measures of criticality-associated phenomena during attention lapses, as measured by slow RTs. Specifically, we calculate several metrics to help generate a better picture of the state of the system (see Fig 2). In particular, we calculated LRTCs to indicate whether there was a deviation from criticality before an attention lapse. Then, we used spectral slope alongside theta (3-7 Hz) and alpha (8-12 Hz) power and variability to help determine the direction of these changes and the overall fit of the edge of synchrony model for brain dynamics and attention. We focused on alpha (8-12 Hz) and theta (3-7 Hz) oscillatory bands, as prior research has consistently identified changes in these bands in ADHD [33, 34]. In combination, these metrics should allow us to establish the direction of deviation relative to criticality, if it exists, during response slowing. For example, if slow reaction times are preceded by shifts toward the asynchronous regime (resulting in increased neural variability or “noise”) and away from criticality, then we would predict (Fig 2), before slow RTs:

1. LRTCs will *decrease* as dynamics shift further towards the asynchronous phase and away from criticality
2. Spectral slopes will get *flatter*

**Figure 2:**
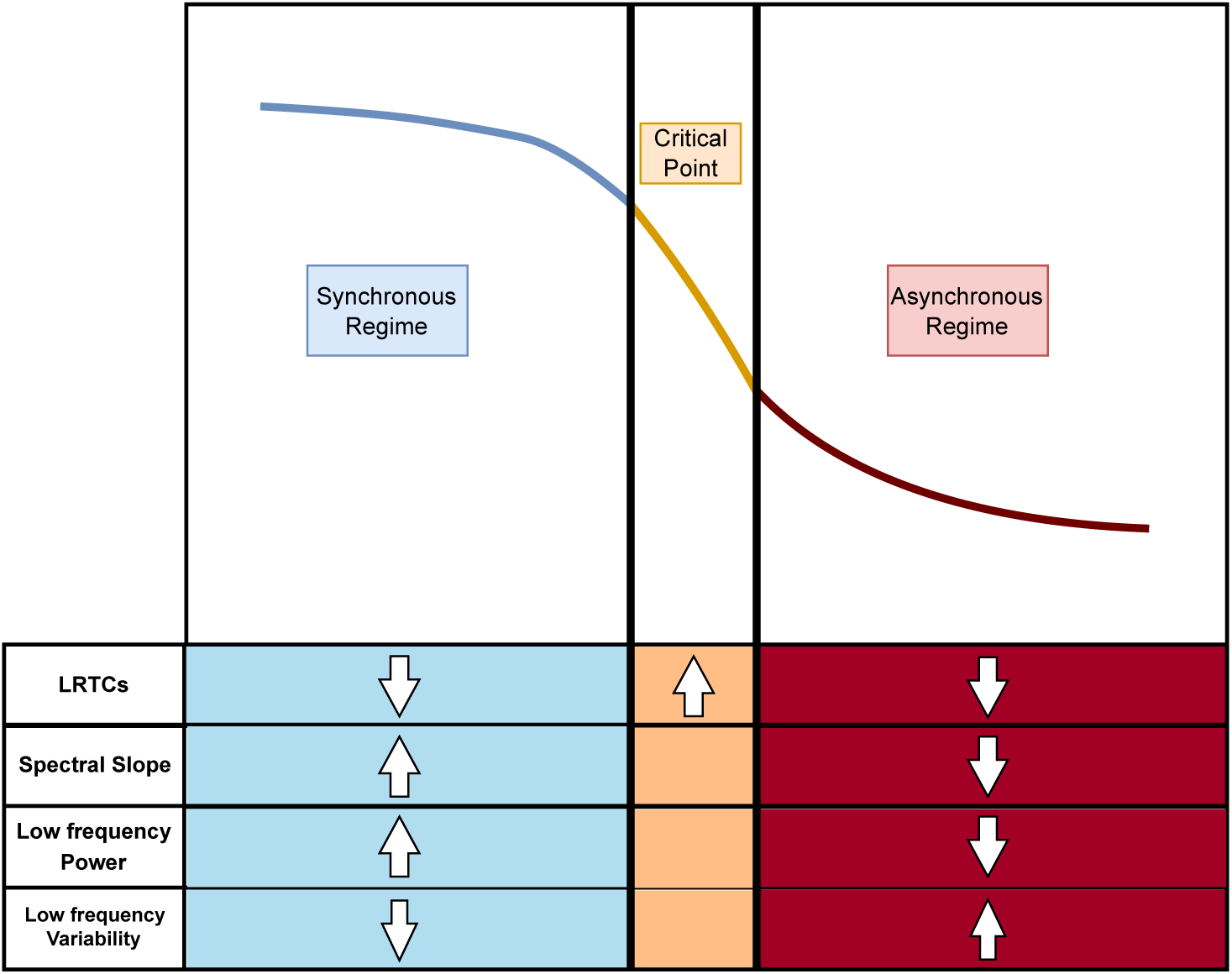
Hypothesized metric changes around edge-of-synchrony criticality. This figure depicts how the direction of the measures should move if the system is moving around the critical point. Crucially, the only metric that indicates distance from criticality are LRTCs. Deviations from criticality towards the synchronous phase should be associated with steepening slopes and flatter slopes for deviations towards the asynchronous phase. Under the edge-of-synchrony model, oscillatory power and variability are expected to shift inversely. Synchronous deviations should increase low-frequency power and reduce variability; while asynchronous deviations do the opposite. Based on prior literature indicating that ADHD and task results in flatter slopes, ADHD individuals should show a similar pattern to typically developing controls where task results in a shift towards the asynchronous regime with attention lapses being instances where dynamics stray too far into this regime. Under this hypothesis, the between-subject effects seen in ADHD are not due necessarily to incorrect within-subject modulation, but due to a difference in “starting” resting state dynamics. As individuals with ADHD are already shifted towards the asynchronous regime at rest, it is more likely that task-related modulations would result in straying too far into the asynchronous regime, leading to increased attention lapses and RTV.

Furthermore, if the dynamics support the presence of edge-of-synchrony criticality we expect that:

1. Low-frequency oscillatory power will *decrease*
2. Low-frequency oscillatory variability will *increase*

Similarly, if differences between individuals with and without ADHD are accounted for by the same criticality-associated neural dynamics as for slow versus fast reaction times, then we also predict that individuals with ADHD will have lower LRTCs, flatter slopes, increased low-frequency variability and decreased power.

## Materials and Methods

### Participants

We analyzed brain activity from a sample of adults diagnosed with ADHD (N=103) and healthy control participants (N=32) previously described in [35, 36] (see Table 1). The recruitment, diagnosis and testing were completed at the University of California, Los Angeles (UCLA). Participants underwent a full clinical diagnostic protocol which included a clinical diagnosis by a clinician (sufficient diagnostic criteria to meet DSM-5 diagnosis, confirmed by evidence for childhood onset ADHD), as well as a battery of supplemental neurocognitive assessment, including the ASRS 18-item, Wender Utah Rating Scale, BRIEF-A, Attention Awareness Questionnaire, Hypersexuality questionnaire, Mood Disorder Questionnaire, HELPS, ACDS, CGI, MINI, WASI-II, WAIS IQ, DK Trails Test, Color word interference task, Five Facet Mindfulness Questionnaire, Sleep Questionnaires, Connors CPT, and Spatial Working Memory Capacity Test. Participants then participated in EEG-data collection during which they performed an audio-visual continuous performance sustained attention task. The study was approved by the UCLA Institutional Review Board. All participants provided informed consent at the time of testing. The Wender Utah Rating Scale, BRIEF-A, Attention Awareness Questionnaire, Hypersexuality questionnaire, Mood Disorder Questionnaire, HELPS, ACDS, CGI, MINI, WASI-II, WAIS IQ, DK Trails Test, Color word interference task, Five Facet Mindfulness Questionnaire, Sleep Questionnaires, Connors CPT, and Spatial Working Memory Capacity Test are not the focus of the current study and will be discussed elsewhere. Participants who were taking medication were instructed to observe a 24-hour washout period before attending their study session. All data were obtained from a previously approved study reviewed and approved by the Medical Review Board at UCLA. All participants gave informed consent in accordance with the approved protocol.

**Table 1:** Participant demographics.

|  | Control (N=32) | ADHD (N=103) |
| --- | --- | --- |
| Age (years) | 25.12 ± 5.7 | 25.47 ± 5.51 |
| Gender (Female) | 54.55% | 60.19% |
| ASRS inattentive | 10.38 ± 5.52 | 26.18 ± 4.25 |
| ASRS hyperactive | 9.32 ± 5.63 | 21.38 ± 5.58 |

### Task

The task design and protocol was identical to [37]. In brief, participants were presented with a stream of auditory tones and Gabor patches and asked to respond to either the auditory or visual stimulus while ignoring the stream of the other modality. Participants completed three blocks, each consisting of six 40 second runs. The runs alternated between three task conditions: 2 passive where both streams were presented but the participant was not asked to respond to either stimulus, 2 auditory-attend where the participants had to attend and respond to the auditory stimuli, and 2-visual attend where participants had to attend and respond to the visual stimuli. During the attend conditions, the participants were asked to make a binary decision for each stimulus to confirm that participants were actively attending to the task. The passive task instructions included a request for participants to fixate on the screen. Video was used to confirm that participants were attending to the screen during the passive viewing task to ensure that participants were paying attention. Each run comprised 25 auditory and 25 visual stimuli, presented in independent streams, and was followed by a 17 second break. The order of the runs was random across blocks and subjects except for the passive viewing runs that were always at the beginning and end of the blocks.

Within each run, the ISI for stimulus presentation was randomly sampled from a uniform distribution ranging from 700 to 2000 msec, separately for each sensory stream. To avoid overlapping presentation of auditory and visual stimuli, the ISIs were further adjusted so that a minimum of 350ms passed between the presentation of one stimulus to the next, resulting in the ISI range of 8502300 msec for stimuli within the same sensory modality and an ISI of 350800 ms between any two stimuli. The auditory pure tones (sampled at 22,050 Hz, 10 msec ramp up and down) and visual gratings were created in Matlab [38]. The experiment was programmed in PsychToolbox [39, 40].

Attention lapses have previously been associated with the exponential, or lengthened tail, of RT distributions that is said to represent “abnormally slow” responses [1]. Meanwhile attention slips, or errors, have been associated with faster responses [41, 42, 43]. Given that both tails of the RT distribution may represent separate but related decreases in sustained attention, we divided each participant’s responses into four event types: fast, average, slow, and passive. Fast responses corresponded to those RTs less than or equal to the 1st quartile (*<*= 25^th^ percentile) and the slow responses greater than or equal to the 3rd quartile (*>*=75^th^ percentile). The average responses were those that fell between the two percentiles, representing the average response time for that individual. We also randomly selected 25 stimuli during passive viewing as a non-task reference that would also control for baseline alpha. To test whether changes in criticality-associated EEG features predict changes in response time, we focused on the 10s preceding the four event types by segmenting each stimulus from -10s to -0.005s, assigning it to one of the four event types based on response time (fast, average, slow) or task (auditory, visual, or passive), and calculated six features: the first cumulant, spectral slope, theta power and variability, and alpha power and variability.

### EEG Recording, Preprocessing, & Analysis

During the task, EEG data were recorded using a 256-electrodeHydroCel Geodesic Sensor Net (EGI, Eugene, OR), digitized using a Net Amps 300 amplifier (10,000 Hz anti-aliasing filter; common-mode rejection 90 dB; input impedance 200*M* Ω), sampled at 500 Hz, and referenced to the vertex electrode (Cz). Electrode impedances were kept below 50*k*Ω.

EEG preprocessing and analysis was done in Python with custom scripts and MNE [44, 45]. The blocks were processed as continuous and cleaned using a combination of artifact subspace reconstruction (ASR) [46] with a cutoff of 20-25, and ICA using the extended Picard algorithm [47]. ICA components associated with eye movement, muscle, and ballistocardiogram were identified using the MNE implementation of ICLabel [29]. It was then bandpass filtered at 1 45 Hz with a bandstop at 60 Hz (zero-phase, hamming window FIR filter), and referenced to the average. The autoreject library was used as a last pass to drop bad trials [48, 49] where 3 channels were the maximum allowed to be interpolated per trial and the threshold set between 0.4 - 0.5 which represents the percentage of channels that must agree.

To reduce dimensionality and increase signal-to-noise ratio, spatial clusters of six electrodes based on Delauney triangulation were created where the centroid of each cluster was on electrodes: F3, Fcz, F4, C3, C1, C2, C4, P3, Pz, P4, O1, Oz, O2 (Figure 3). Each metric was calculated for each electrode of the cluster and then averaged together to get the ROI value. If an electrode of the cluster was dropped due to excess noise, the whole cluster was skipped as interpolation could already have been applied during the autorejection step and a second interpolation could result in excess loss of dimensionality. For the auditory task, the response at each electrode cluster was then averaged across the scalp while only the occipital clusters were averaged together for the visual task due to the spatial specificity of the neural response to visual tasks.

**Figure 3:**
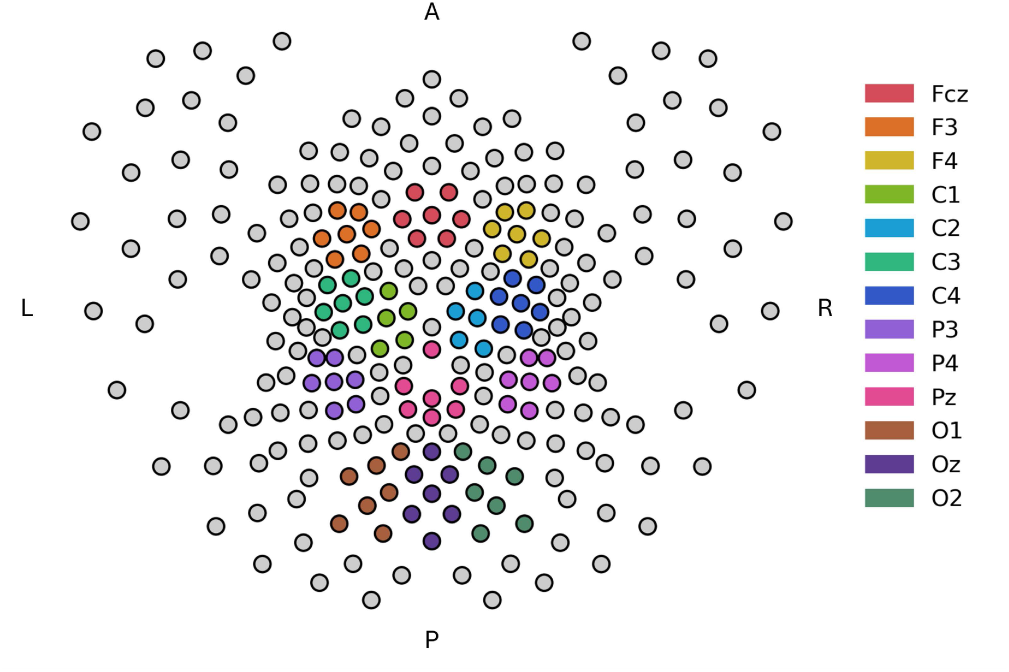
ROI clusters. The colors of the figure indicate the electrode clusters for which the measures were calculated. Due to the high density of the cap, neighboring electrodes have very similar signals, so to reduce dimensionality of the dataset, these clusters were identified by Delauney triangulation, the mean of the channels taken, and each EEG metric was applied to this mean of the cluster.

### Feature Extraction

We computed the first cumulant using the Wavelet leader multifractal analysis (WLMA; [50, 51]). The WLMA can be used to assess whether a signal is monofractal (the distribution has a small width) or multifractal (there is significant distribution of behavior at different scales) and describe the dominant scaling behavior (i.e., statistical characteristics) and its variability across timescales. In other words, it assesses the local regularity or scaling of a signal, quantifying both the dominant scaling behavior (the first cumulant) and the distribution or variability of this behavior (higher order cumulants). In our data, the vast majority of trials across individuals were not multifractal, so the analysis focused on the first cumulant. The first cumulant was calculated using the WLMA and wavelet coefficients with Debauchies wavelets with *N_ω_*=1 vanishing moments and over the octave ranges of 2 to 7 (associated with frequencies 1 - 30 Hz), implemented in the pymultifracs toolbox (https://github.com/neurospin/pymultifracs) (for a discussion on the window size, see Appendix C). A decrease in the cumulant is indicative of a deviation from criticality, but does not indicate towards which of the two phases (synchronous or asynchronous). Spectral slope was calculated using the same discrete wavelet transform (DWT) from the WLMA and FOOOF. As the slope results were more consistent across tasks for the DWT fit, we focus on those results, but the FOOOF results are also reported (see Appendix A). Unlike the cumulant, the direction of change in the spectral slope indicates the direction of the deviation where a steeper slope indicates a shift towards the synchronous phase and a flattening a shift towards the asynchronous phase.

The power of theta (4 7 Hz) and alpha (8 12 Hz) oscillations over and above the aperiodic component (spectral slope) was calculated using specparam. To measure the variability in the power of both alpha and theta, a time-frequency decomposition was calculated using Morlet wavelets with 3 cycles and linearly spaced between 8 12 (alpha) and 4 7 (theta) and then the coefficient of variation was calculated for each epoch. Proximity to criticality has been shown to affect peak frequency and power in alpha [52], such that a shift toward the synchronous phase would be associated with increased power with opposite effects in asynchronous phase. We also include amplitude variability as a more overt metric of variability connected to oscillations. Decreased amplitude variability of oscillations should coincide with increased power as when the system transitions into a synchronous regime, coupled oscillators begin to align in phase and oscillate collectively. This alignment reduces the random, conflicting perturbations exchanged between oscillators, allowing their amplitudes to stabilize. Thus, we hypothesize that the amplitude variability in these bands should decrease as the system transitions to a fully synchronous state and the oscillatory power increases.

### Statistical Analyses

We were interested in assessing both the within and between-subject effects in RT variability and thus intra- and inter-individual variability. A 2 (diagnosis) x 2 (task) mixed ANOVA was run for mean RT, RT standard deviation (RT STD), and RT coefficient of variation (RT CV) and percent correct in JASP [53] with both gender and age as covariates. Age was a significant covariate for the mean RT and RT STD, but not RT CV. Because RT CV is a normalized version of RT STD by each individuals mean RT, it likely diminishes the differences due to age on the metric. For the EEG metrics, a 2 (diagnosis) x 4 (event type) mixed ANOVA was run for each measure in JASP and were split by modality (Auditory and Visual) for robustness. Both gender and age were added as covariates to all analyses, but only the age covariate remained significant for the wavelet- and multitaper-based slopes (consistent with previous literature indicating a strong association between spectral slope and age [9]), so gender and age were dropped for the remaining analyses. The Greenhouse-Geisser sphericity correction was applied when Mauchlys test indicated that the assumption of sphericity was violated. Due to the unequal sample sizes, we also did an assumption check for homogeneity using Levene’s test. Due to a violation of the assumption of homogeneity of variance for theta and alpha CV for both auditory and visual tasks, a stratified non-parametric permutation with 1,000 iterations was used to calculate robust p-values using the afex, boot, and boot.pval packages in R [54]. The Holm correction was used for post-hoc comparisons. We provide exact p-values when p is greater than 0.001. Between-subject variability is thus assessed by the main effect of diagnosis and within-subject effects by task. The alpha level was set to 0.05.

## Results

### Behavioral task and RTV

In general, the auditory task showed more variable performance than the visual task for both groups (main effect of task for RT CV: *F* (1, 132) = 8.04*, p* = 0.005), but this variability did not result in a significant decrease in performance between task modalities. All subjects performed well with a mean accuracy of 94% across groups and task, but individuals with ADHD were less accurate overall than the control group (ADHD=93.3% and control group=97%: *F* (1, 132) = 26.22*, p <* 0.001). Along with being less accurate, individuals with ADHD took longer on average to respond (main effect of diagnosis for mean RT: *F* (1, 132) = 4.11*, p* = 0.045) and were more variable in their performance (main effect of diagnosis for RTCV and RT STD: *F* (1, 132) = 6.80*, p* = 0.017; *F* (1, 132) = 5.84*, p* = 0.01). Thus, as expected, participants with ADHD showed increased RTV and decreased accuracy during sustained attention across visual and auditory modalities (see Table 2).

**Table 2:** Task performance (mean and standard deviation) split by modality and diagnosis. Accuracy is reported as percent correct while RT CV denotes the coefficient of variation of reaction time. Significant diagnostic effects are shown for each modality by an asterisk on the ADHD values (* p < .05, ** p < .01, *** p < .001). *Note:* For accuracy, the asterisk represents the significant results from a permutation test due to the violation of the as<u>sumption of homogeneity of variance.</u>.

| Metric | Auditory |  | Visual |  |
| --- | --- | --- | --- | --- |
|  | Control (N=32) | ADHD (N=103) | Control (N=32) | ADHD (N=103) |
| Accuracy (%) | 96.3 ± 0.03 | 93.4 ± 0.04*** | 96.8 ± 0.02 | 93.1 ± 0.03*** |
| Mean RT (ms) | 493.91 ± 52.32 | 514.04 ± 62.60* | 489.21 ± 39.22 | 509.45 ± 49.66* |
| RT SD (ms) | 128 ± 39 | 148 ± 41** | 98 ± 26 | 114 ± 33** |
| RT CV (%) | 25.56 ± 6.11 | 28.50 ± 5.90* | 19.70 ± 4.54 | 22.14 ± 5.37* |

### Slowing reflects a shift away from asynchronous regime during task

We first investigated within-subject dynamics, namely how LRTCs and spectral slopes change during task and before slow RTs. The first cumulant showed a main effect of event type in the auditory task (*F* (1.66, 226.3) = 3.262*, p* = 0.049) and the visual task (*F* (1.68, 221.20) = 5.869*, p* = 0.005). Post-hoc comparisons showed that this effect was due to a significant *increase* in LRTCs preceding slow responses compared to fast and average responses (*p* = 0.039) in the auditory task (Fig. 4). Similarly, in the visual task LRTCs increased prior to slow response and during passive viewing compared to periods preceding average response times (average vs slow *p* = 0.015 and average vs passive *p* = 0.007). These findings suggest that slow responses during task are preceded by an *increase* in LRTCs, which would be consistent with a shift *towards* criticality, and potentially away from the task-appropriate state. To examine whether this was asynchronous or synchronous regime, we evaluated spectral slopes, which also showed significant effects of event type. During the auditory task, the spectral slopes showed a main effect of event type (*F* (1.63, 220.59) = 5.45*, p* = 0.008), with post-hoc comparisons indicating that passive viewing and slow responses were preceded by *steeper* slopes than fast (*p* = 0.025 and *p* = 0.014) and average task responses (*p* = 0.016 and *p* = 0.025), which according to our model would suggest a shift away from the asynchronous regime. In combination, the LRTCs and slope effects suggest that task-state is asynchronous and RT slowing (as well as passive events) are accompanied by a shift towards the critical point (moving right to left in Figure 2). This finding is notable as, while consistent with prior literature suggesting that the task state compared to resting or a passive viewing state involves a deviation from criticality, contrary to expectations assuming a noise hypothesis of RT slowing, we found that dynamics shifted away from the asynchronous regime and towards criticality prior to response slowing (Fig. 4).

**Figure 4:**
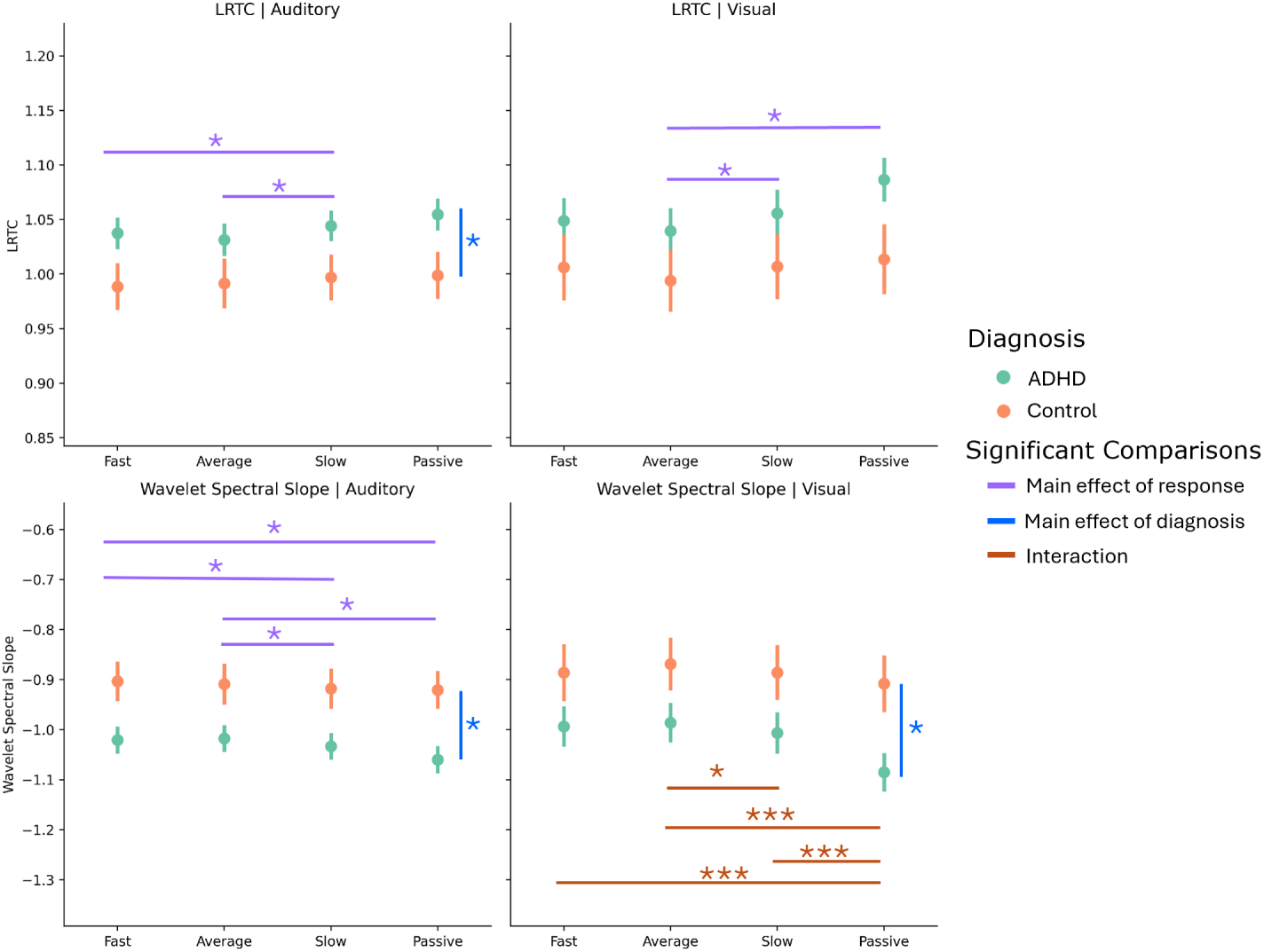
Slow responses are preceded by steeper slopes and more LRTCs. Both wavelet-based slope and LRTCs show differences between the fast and average responses compared to the slow and passive control conditions where the slow responses appear are similar to the passive condition. This effect appears primarily driven by the ADHD group, except for the wavelet-based slope during the visual task where only ADHD shows this pattern. *Note:* The colors of the bars indicate whether the effect is due to an interaction (orange) or a main effect of response (purple) or diagnosis (blue). The asterisks above the bars denote significance (* *p <* 0.05, ** *p <* 0.001, and *** *p <* 0.001). Error bars represent the standard error.

### Individual differences in criticality dynamics echo slowing effects

In examining between-subject effects, we found effects of diagnosis that were analogous to those of RT slowing. Namely, LRTCs showed a main effect of diagnosis for the auditory task where the ADHD group was associated with *increased* LRTCs compared to the control group (*F* (1, 136) = 3.923*, p* = 0.05), consistent with a state in ADHD closer to criticality. The diagnosis effect was not significant however in the visual task, consistent with lower behavioral variability in this task. For the spectral slope, once again there was a significant main effect of diagnosis, where the ADHD group had significantly *steeper* slopes than the control group (*F* (1, 135) = 7.384*, p* = 0.007), echoing the steepening of slopes for slower RTs. However, in the visual task, the spectral slope also showed a significant interaction effect between the type of event and the diagnosis (*F* (1.63, 213.00) = 3.37*, p* = 0.046). This was because the effect of the event on the slope (i.e., a steeper slope before passive viewing compared to fast and average responses and slow RT compared to average responses) (passive: *p <* 0.001 and average vs slow: *p* = 0.033), was only present in the ADHD group. So, while the steep slope pattern before RT slowing and passive viewing is similar between auditory and visual tasks, the control group showed this pattern only during auditory tasks while the ADHD group showed it during both tasks (Fig 4).

Taken together, these results suggest that prior to RT slowing, brain dynamics shift away from the asynchronous state, towards criticality. Given that similar effects were observed for passive trials, this also suggest that RT slowing approximates a return towards a resting or baseline state. The interactions with group, further suggests that this pattern is more consistent across tasks in ADHD. Importantly, the combined group effects suggest that these dynamics are exaggerated in ADHD, consistent with a baseline or “starting” point closer to criticality. This contradicts the hypothesis that more variable task performance reflects an excessive shift towards the asynchronous regime. Rather, it suggests that more variable task performance in continuous performance tasks is related to insufficient shifting towards asynchrony.

### Oscillatory dynamics diverge from edge-of-synchrony predictions

To clarify whether the edge-of-synchrony model is the most appropriate to explain the between-subject effects and within-subject dynamics, we next analyzed low-frequency power and variability. If we extend this unexpected direction towards a more critical regime prior to RT slowing, then, according to the edge-of-synchrony model, we would expect that low-frequency oscillations would show *increased* power and *decreased* variability (see, Fig. 2). However, neither low-frequency power nor variability followed these predictions.

Low-frequency power primarily reflected stable between-subject differences rather than dynamic task-related changes before RT slowing. There were no significant findings for either alpha power during the auditory task, but in the visual task, alpha power did show a main interaction effect (*F* (1.81, 238.22) = 4.15, *p* = 0.02) and main effect of response (*F* (1.81, 238.22) = 3.77*, p* = 0.029), but no post-hoc comparisons were significant for either effects. On the other hand, theta power showed a main effect of diagnosis in the auditory task (*F* (1, 135) = 4.86*, p* = 0.029) and visual task (*F* (1, 131) = 4.82*, p* = 0.03) with the ADHD group showing higher theta power overall compared to control group. During the visual task, there was also a main interaction effect between event type and diagnosis (*F* (1.95, 255.77) = 3.61*, p* = 0.029) with post-hoc comparisons showing that theta power increased during passive viewing compared to task (across all other event types), but only in ADHD (all *p <* 0.001) (Fig. 5). These results indicate that between-subject theta power during passive viewing distinguishes between adults with and without ADHD, but are not predictive of behavioral performance.

**Figure 5:**
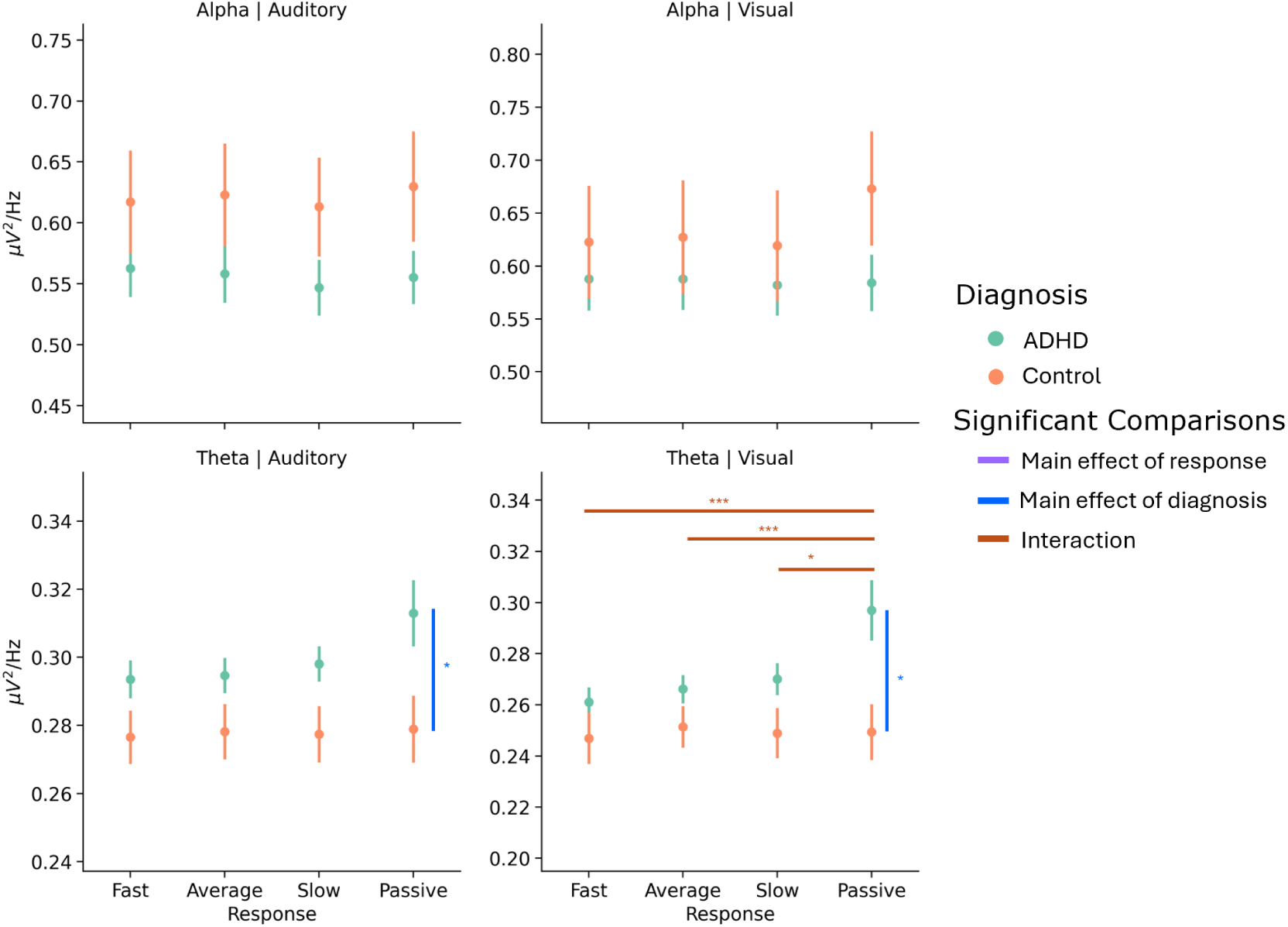
LF power mainly follows group differences at rest. Theta power was elevated during rest in ADHD compared to controls, but decreases during task. This difference implies that the dynamics that control or modulate theta power are similar during task between ADHD and controls, but diverge during rest. *Note:* The colors of the bars indicate whether the effect is due to an interaction (orange) or a main effect of response (purple) or diagnosis (blue). The asterisks above the bars denote significance (* *p <* 0.05, ** *p <* 0.001, and *** *p <* 0.001). Error bars represent the standard error.

Low-frequency variability, instead, showed task-related changes before RT slowing, but variability increased only in the ADHD group. Namely, during the auditory task, there was an interaction between diagnosis and event type (alpha: *F* (1.969, 267.79) = 7.91*, p_perm_* = 0.001, theta: *F* (1.634, 222.24) = 3.47*, p_perm_ <* 0.001), along with a significant main effect of event type (theta: *F* (1.63, 222.24) = 22.75*, p <* 0.001, alpha: *F* (1.969, 267.79) = 11.58*, p <* 0.001) and diagnostic category (theta: *F* (1, 136) = 6.496*, p* = 0.012). Post-hoc comparisons for both theta and alpha CV indicated that neural oscillations preceding slow RTs were more variable than those preceding fast and average responses, but only for the ADHD group (all *p <* 0.001) (Fig. 6). Both ADHD and controls showed significant differences between passive viewing and fast and average responses (ADHD: *p <* 0.001, Controls: *p* = 0.02). The main effect of diagnosis for theta indicated that individuals with ADHD had higher theta variability overall compared to the control group. The effects were analogous in the visual task. The interaction between diagnosis and event type was significant at trend-level (theta: *p_perm_* = 0.07, alpha: *p_perm_* = 0.065), but the main effects of event type were significant for both theta (*F* (1.65, 217.37) = 14.22*, p <* 0.001) and alpha CV (*F* (1.78, 235.38) = 10.19*, p <* 0.001). For theta CV, variability of neural oscillations was higher preceding passive viewing compared to average responses (*p* = 0.032), and slow responses compared to fast (*p <* 0.001) and average responses (*p <* 0.001). The same pattern was present for alpha CV (average and fast v passive, both *p <* 0.001 and average v slow, *p* = 0.038) but fast was also significantly different to slow (*p* = 0.007) (Fig. 6). Similarly, the main effect of diagnosis (theta: *F* (1, 132) = 6.07*, p_perm_* = 0.013 and alpha: *F* (1, 132) = 5.22*, p_perm_* = 0.028) indicated that individuals with ADHD had higher alpha and theta variability overall across passive viewing and the visual task compared to the control group.

**Figure 6:**
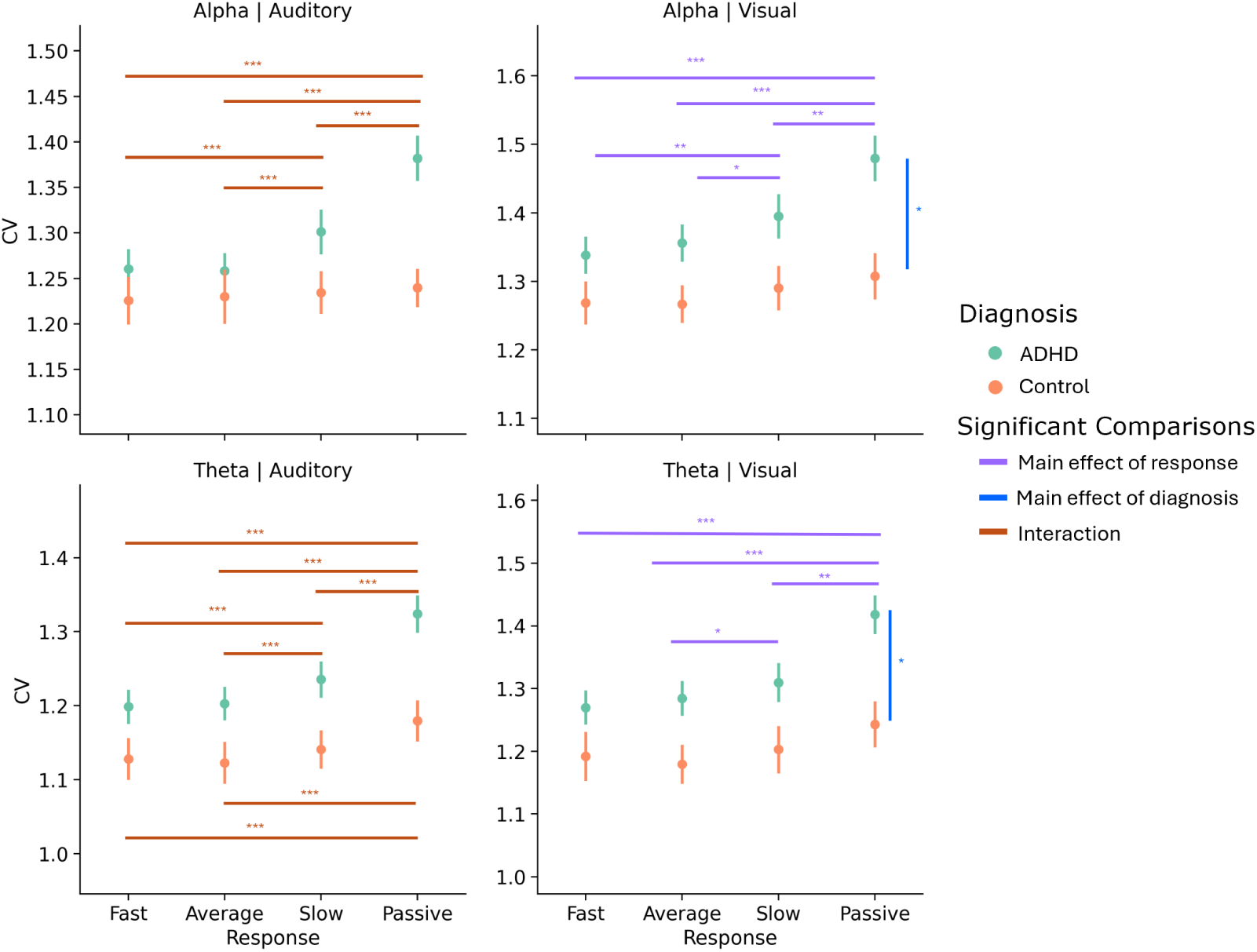
LF Variability in ADHD shows a similar pattern to LRTCs and spectral slopes. This pattern described by LRTCs and spectral slopes is present for LF variability, but is dependent on diagnosis and task. For the auditory task, only the ADHD show this pattern (left column) while for the visual task both controls and ADHD show it (right column). Furthermore, during the visual task, the ADHD group still shows increased LF variability compared to controls while for the auditory task, the difference is only noticeable at rest and appears to be diminished during task. *Note:* The colors of the bars indicate whether the effect is due to an interaction (orange) or a main effect of response (purple) or diagnosis (blue). The asterisks above the bars denote significance (* *p <* 0.05, ** *p <* 0.001, and *** *p <* 0.001). Error bars represent the standard error.

In summary, low-frequency power and variability is indicative of ADHD-specific oscillatory dynamics that are not consistent with edge-of-synchrony criticality. However, as the pattern of variability change during task is the same as the one seen for LRTCs and spectral slopes, the results are generally consistent with criticality, and imply that perhaps another critical point model may better fit the data.

## Discussion

In this study we investigated whether RT slowing, a key contributor to RTV and associated with attention lapses can be predicted by changes in criticality, and whether between-subject differences between adults with and without ADHD showed similar differences. The results replicate prior findings of a shift in dynamics during task away from criticality relative to resting state [55, 30, 56]. However, we found that slow RTs were preceded by a shift back towards criticality, resembling the dynamics observed during a passive non-task condition, and not further into the asynchronous regime as hypothesized given the noise hypothesis (Fig. 2). ADHD mirrored these differences at the group level where across both task and rest, individuals with ADHD were closer to criticality than controls. However, while we do find evidence consistent with criticality in adults with and without ADHD, changes in low-frequency variability mainly in adults wtih ADHD were not consistent with edge-of-synchrony criticality, suggesting that a different critical point may be involved in this population. Finally, we find that the between-subject differences in low frequency power are not related to within-subject changes in neural dynamics or RT which suggests that differences in power likely reflect more general differences in neural state.

Our results are consistent with the functional interpretation of the “noise” hypothesis of ADHD, namely that individuals with ADHD exhibit neural dynamics that are consistent with more “exploratory” behavior typically associated with criticality. Yet, our results suggest that this exploratory behavior is not “noise,” defined as random variability. Rather, our results indicate that prior to slowing behavioral performance, and in ADHD, dynamics are closer to criticality (shifting away from the asynchronous regime). This indicates that although there is increased neural variability, it is more *structured*. By structure, we are referring to the increased LRTCs, which signal that the statistical characteristics are self-similar or similar across scales and thus more flexible and robust. For this reason, proximity to criticality has been associated with increased cognitive flexibility [57] and cognitive effort [58]. Indeed, it has been suggested that slow RTs represent moments when individuals “catch” themselves before a full lapse (resulting in an error) occurs [2]. The dynamics of slow RTs observed here may reflect such a process of reorienting and the increased proximity to criticality in participants with ADHD may suggest that the re-orienting or “catching” was more effortful. An alternate possibility, consistent with our data, is that the periods prior to RT slowing reflect mind wandering. The proximity to criticality is consistent with a drift in attention to more internal task-unrelated thoughts–similar to the phenomenological experience of rest. This highlights how important perceptions of task difficulty and strategies are to understanding the brain dynamics that underlie RTV. In the first case of “catching,” increased LRTCs may be indicative of better integration of information which is incompatible with the “noise” hypothesis. While in the case of mind wandering, increased LRTCs are more inline with the functional interpretation of the “noise” hypothesis. As the current paradigm is unable to disentangle whether the shift towards criticality before slow RTs reflects shifts to task-unrelated thoughts or task-related processes, subsequent studies should focus on connecting the phenomenological experiences of attention lapses, slips, and task-related interference with shifts towards criticality.

Collectively, our results suggest that the mechanisms behind within-task changes in sustained attention appear to remain relatively intact in adults with ADHD with the main differences in neural dynamics resulting from a difference in “starting” dynamics at rest that carry down into task. The neural dynamics in adults with ADHD started and remained closer to criticality through task and passive viewing. This increased proximity to criticality may increase the chances that a random fluctuation in dynamics pushes the system too close to criticality, resulting in more instances of increased cognitive effort to re-orient or increased task-unrelated thoughts in individuals with ADHD and bigger changes in dynamics. These bigger changes and increased incidences would also explain why the ADHD group also showed more consistency in the within-subject event effects (where dynamics before RT slowing were similar to those during passive trials) across modalities and features. Conceptually, this resembles the existence of different attractors or set points in the system’s state space (referring to an N-dimensional space that contains all possible states or parameter configurations). In this framework, adults with ADHD tend to stabilize near a set point that lies closer to the system’s critical threshold. However, the starting dynamics do not seem to be the whole story, as when we investigated whether the EEG metrics before slow RTs and at rest correlated with overall behavioral performance (mean RT, RT standard deviation and correlation of variation, and precision) or symptom severity, no correlations survived corrections for FWER (see Appendix B). We recognize that, it is possible that adults without ADHD may exhibit the same dynamics as adults with ADHD (i.e., principally, the interaction results: LF variability, power, and spectral slope), but due to the low sample size of adults without ADHD there may not be enough power to detect the pattern in that group. Future work is needed to investigate the precise association between resting state dynamics and their effect on task dynamics and sensory processing along with overall behavioral performance to better understand how differences in “starting” dynamics in ADHD are related to RTV.

Finally, we address low frequency power and variability which did not follow the predicted hypothesis regarding the role of edge-of-synchrony in criticality dynamics during behavioral slowing. For LF power, only theta power showed between-subject differences at rest between individuals with ADHD and the control group which aligns with previous work [59, 60, 33, 61, 62, 63], but not with predicted changes within-individual. Thus, low frequency power may reflect differences more broadly in arousal or motivation indicative of the general dynamic phase space that each individual occupies at the moment of recording. While LF variability showed a qualitatively similar pattern to LRTCs and spectral slope (dynamics preceding slow RTs were similar to those during rest), in ADHD, it *increased* instead of decreasing (as would be suggested by the steepening of the slope). As such these results are not consistent with the hypothesized connection between spectral slope and oscillatory amplitudes in an edge-of-synchrony regime. Principally, we found that spectral slope *steepened* in the same periods when variability *increased*. Neural models have previously tied flattening of the spectral slope to an increase in excitation, which suggests more variability [64]. This result is, therefore, inconsistent with the simple edge-of-synchrony model that we proposed. However, the results still bear hallmarks of a critical system. Namely, criticality can be identified either by the rapid increase in particular phenomena or their maximization (i.e., LRTCs) and for other types of criticality, variability is not expected to increase monotonically (like for edge-of-synchrony) but to be maximized at criticality (e.g., avalanche criticality). The increase in LF variability at rest and prior to RT slowing primarily in ADHD may be due to their increased proximity to some other critical point compared to adults without ADHD, possibly one where variability is maximized. In other words, the phase space in ADHD may have a different dynamic profile with different type(s) of critical points. Thus, although the current experimental data is not fully consistent with the edge-of-synchrony, it does still suggest that criticality is involved in some way. These findings also underscore the importance of measuring a variety of criticality-associated phenomena, when trying to make inferences about underlying neural dynamics. Without additional measures of oscillatory power and variability, the nuanced differences in neural activity between ADHD and control groups might have been overlooked. A more limited framework that relied solely on LRTCs or slopes alone would have lost the nuance given by the slope and LF variability. These findings highlight the importance of approaching criticality from a strong theoretical foundationgrounded in a specific critical pointso that empirical observations can be meaningfully interpreted within the appropriate framework.

Finally, it is notable that the complex relationship between within- and between-subject effects would have been missed if we relied solely on between-subject differences, emphasizing the importance of fully evaluating both within and between-subject variance. As illustrated by the discussion above, consideration of both effects is important in determining which hypotheses or theories of neural mechanisms best predict and describe cognitive processes (for a more thorough discussion of the intricacies of this relationship, please see [65]). Consideration of both the within-subject and between-subject effects allowed us to identify more clearly the differences and similarities of the neural mechanisms that underlie decreases in behavioral performance between ADHD and controls. Together our results suggest that while low frequency power can distinguish between ADHD and controls, LRTCs, spectral slope, and LF variability are better predictors of differences in performance between and within the groups. Low frequency power may reflect more broadly differences in arousal or motivation indicative of the general dynamic phase space that each individual occupies at the moment of recording, and which affects their general abilities, but it is the subtle changes in LRTCs, spectral slope, and LF variability, and thus system dynamics, that predicts behavioral performance. By looking at both within- and between-subject effects and multiple overlapping phenomena associated with criticality, we were able to find evidence that while the edge-of-synchrony criticality does not fully fit the described behavior, brain dynamics prior to RT slowing show signs of a shift towards some critical point, indicative of more “exploratory” or computationally intensive behavior and not more “random” noise.

In conclusion, our findings implicate a relationship between critical brain dynamics and behavioral performance. Specifically, slower response times were consistently preceded by more “statistically structured” neural variability, a signature of increased proximity to criticality, and were not consistent with a simple “noise” interpretation. This pattern held across adults with and without ADHD, although individuals with ADHD tended to be generally closer to critical-itypotentially contributing to their increased RTV. Notably, the observed dynamics diverged from the hypothesized edge-of-synchrony model, suggesting the presence of a different critical regime. More broadly, these results highlight the importance of integrating within- and between-subject variability when probing how different dynamical states and models relate to behavior and neuropsychiatric conditions.

## Acknowledgments

This work was supported by funding from the National Institutes of Health to AL (MH116268, MH128475, MH130731) and GVS (R44MH099709, R43MH101924) and from the German Research Foundation (ZI 1895/1-1) awarded to NZ. We also wish to acknowledge the late Robert Bilder for his contributions to the conceptual design and clinical oversight of the original study from which the present dataset was derived.

## Competing Interests

AL has received payments as a consultant and holds shares in ThinkNowInc. GVS is an officer in Think Now, Inc. and holds shares in the company. All other authors report no biomedical financial interests or potential conflicts of interest.

## Notes

### Summary of Updates

Figure legends revised to include the significant comparison colors. Introduction has been majorly revised to include extended hypotheses. Added new addendum on the chosen window for the LRTC analyses

