## Appendices for "Beyond Neural Noise: Critical Dynamics Predict Slower Reaction Times in Adults With and Without ADHD"

#### Appendix A. FOOOF Results

The spectral slope was also determined using spectral parameterization or the specparam (formerly known as "Fitting Oscillations and One-Over-f" or FOOOF) toolbox [66]. The power spectral density was first calculated for each trial using the multitaper method and then the slope was fit with specparam using the following parameter settings: peak_width_limits = [2,7], max_n_peaks = 5, min_peak_height = 0.05, peak_threshold = 0.2, aperiodic_mode = ‘fixed’.

During the auditory task, the spectral slopes showed a pattern consistent with the LRTCs, with a main effect of event type ($F(1.97, 229.27) = 7.60, p = 0.001$). None of the post hoc comparisons were significant, but the pairwise comparisons between fast vs slow and passive viewing, and average vs passive viewing were trend-level ($p < 0.1$). During the auditory task, spectral slopes also showed a main effect of diagnosis ($F(1, 135) = 4.943, p = 0.028$) where individuals with ADHD had steeper slopes than the control group overall. During the visual task, spectral slopes also showed a significant main effect of diagnosis during the visual task in the occipital electrodes ($F(1, 131) = 4.79, p = 0.043$), indicating once again that ADHD was associated with steeper slopes than control group overall (Fig A.7).

The differences between the FOOOF and Wavelet based slopes is likely due to signal-to-noise (SNR) and, for FOOOF, model fit. Slopes are calculated on every trial instead of the mean PSD of multiple trials which decrease the SNR which in turn affects the model fit for the FOOOF algorithm as it tries to distinguish between periodic and aperiodic behavior. Meanwhile, the discrete wavelet method (DWT) used includes a de-noising step that may help diminish the number of sharp transients and emphasize the broadband scaling trend. The choice between the use of either DWT or FOOOF for calculation of the spectral slope ultimately comes down to whether there is an interest in looking at both the slope and oscillations and the SNR of the underlying signal.

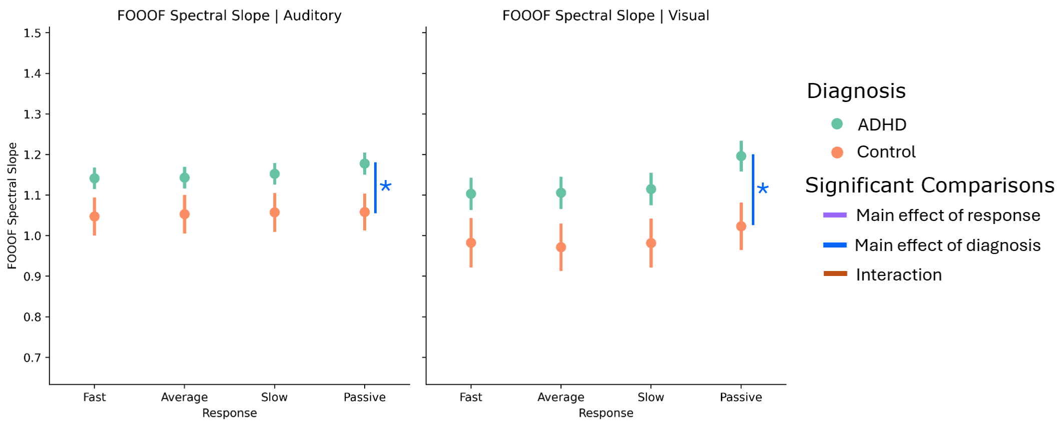


Figure A.7: **FOOOF Slopes are steeper in ADHD at rest.** *Note:* The colors of the bars indicate whether the effect is due to an interaction (orange) or a main effect of response (purple) or diagnosis (blue). The asterisks above the bars denote significance (* $p < 0.05$, ** $p < 0.001$, and *** $p < 0.001$). Error bars represent the standard error.

### Appendix B. Criticality-associated EEG features were not associated with individual differences.

Given the relationship between the calculated EEG features and RT slowing was mirrored by the differences between ADHD and the control group (even though they were in the opposite direction as expected), we lastly looked to see if this pattern of results extended to other individual differences in behavioral performance and ADHD symptoms. We also wanted to compare the results of these correlations with those between a well-known metric in the ADHD literature that has previously been used as a diagnostic aid: the theta/beta ratio (TBR). To do so, we ran Spearmans rank correlation analyses between the z-scored EEG metrics for each response type (fast, average, slow and passive), five performance metrics (mean RT, median RT, RT standard deviation, hit rate, and percent correct), and the ASRS hyperactive and inattentive subscale scores. While spectral slope and LRTCs did not show any significant correlations, the oscillatory metrics (low-frequency variability and power and TBR) did appear to correlate with hyperactivity symptoms and behavioral accuracy. However, none of these correlations survived a subsequent correction for family-wise error.

#### Appendix C. Stability of the WLMA

Due to the short duration of each run (40 seconds for each passive, auditory, and visual run), we choose a smaller window (10s) that is typical for monofractal analysis to minimize overlap between windows. The WLMA was originally created to increase the stability of multifractal measures (like the detrended fluctuation analysis) over shorter windows and signal lengths, but to empirically test that our 10-second windows (sampled at 250 Hz, yielding $N = 2,500$ discrete data points) provide sufficient scale depth for reliable estimation, we looked at the stability of the metric across three longer signal lengths. Specifically, we recalculated the first cumulant for each subject across three passive runs using increasing window sizes of 10, 20, 30, and 40 seconds. Unlike DFA which relies on polynomial detrending of raw signal, the WLMFA leverages mother wavelets and their coefficients to estimate fractality of a signal. The LRTCs are derived based on the wavelet-based coefficients scale range with the number of available wavelet coefficients at a given scale $j$ defined by $N_j = T \cdot f_j$, where $T$ is the window length and $f_j$ is the scale-equivalent frequency. While the coarsest scale (1 Hz) yields 10 coefficients within a 10s window, the finer scales (up to 30 Hz) provide up to 300 coefficients. The monofractal behavior (LRTCs) are then calculated based on the linear relationship between the log-cumulants of these wavelet measures and the range of scales ($j$). Because the final metric is evaluated as a global slope across the entire range, these high-density finer scales help to minimize the impact of the variance at the lowest frequency point. We can see how increasing the number of avaible coefficents at the lowest scale affects this estimation by increasing the window to 40 seconds where the number of available coefficients quadruples from 10 to 40.

The mean within-subject difference in the calculated cumulant, when comparing the 10-second baseline to the 20, 30, and 40-second windows, was 0.015, 0.020, and 0.025, respectively

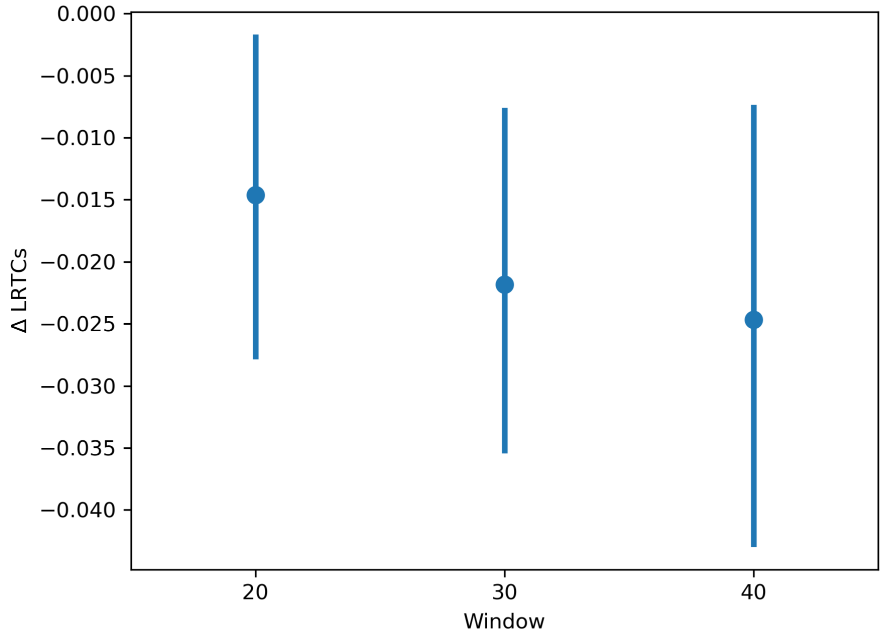


Figure C.8: **Differences in calculated LRTCs based on window size**. The change in the calculated LRTCs between the experimental 10s windows and larger 20, 30, and 40s windows is shown. On average, across subjects, increasing the window size decreases the calculated LRTCs by 0.02. The error bars represent the 95% Confidence Interval.

(Fig. C.8). Furthermore, at the between-subject level, the 95% confidence interval of this mean difference spans from -0.040 to -0.005. While we observed a slight widening of the confidence interval for the 40-second windows, this effect was likely driven by necessary artifact rejection where fewer subjects possessed a continuous, artifact-free 40-second segment compared to the 10-second segments. Because the global regression slope is robustly anchored by the high density of coefficients at the finer scales, the 10-second window effectively calculates the monofractal behavior with negligible variance (a maximum expected shift of only $\sim 0.04$ compared to the longest viable windows).
